# Differential Effects of Short-term and Long-term Deep Brain Stimulation on Striatal Neuronal Excitability in a Dystonia Animal Model

**DOI:** 10.1101/2025.04.07.647541

**Authors:** Marco Heerdegen, Denise Franz, Valentin Neubert, Fabiana Santana-Kragelund, Tina Sellmann, Christoph Werner-Schmolling, Jens Starke, Konstantinos Spiliotis, Franziska Richter, Angelika Richter, Rüdiger Köhling

**Author notes:** corresponding authors Prof. Dr. Rüdiger Köhling, Dr. Marco Heerdegen, Oscar-Langendorff-Institute of Physiology, Rostock University Medical Center, Gertrudenstraße 9, 18057 Rostock, Germany.

## Abstract

Deep brain stimulation (DBS) is, by now, one of the standard treatment options for movement disorders like dystonia or Parkinson’s Disease. Although its clinical effectiveness is established, the exact mechanisms by which DBS influences neural motor networks are not fully understood. The present study explores the development of adaptive network mechanisms with DBS in the dt^sz^ hamster model, an *in-vivo* model exhibiting spontaneous dystonic episodes, by comparing functional impacts of short-term and long-term DBS on medium spiny neurons (MSNs) and synaptic transmission in the striatum. In this electrophysiological study, we uncovered contrasting changes in neuronal excitability and synaptic dynamics following short-term versus long-term DBS. Short-term DBS enhanced neuronal firing responses, while long-term DBS diminished them. Regarding synaptic alterations, both short-term and long-term DBS significantly shifted spontaneous EPSC occurrences to longer intervals, with this effect, however, being more pronounced in short-term DBS, leading to a significant decrease in mEPSC frequency. Notably, acetylcholine application effectively reversed this effect, restoring mEPSC frequency more efficiently again in tissue subjected to short-term DBS compared to long-term DBS. These observations indicate that the therapeutic benefits of DBS in dystonia may involve both immediate and adaptive mechanisms, which has implications for improving stimulation parameters and treatment protocols. The findings shed light on the temporal specificity of DBS effects and highlight the importance of understanding synaptic mechanisms to enhance therapeutic outcomes for dystonia patients.

## Introduction

Dystonia is a neurological condition affecting movement, characterized by persistent or intermittent muscle contractions resulting in abnormal, often repetitive movements or postures (Batla, 2018; Defazio et al., 2013). It is the third most prevalent movement disorder after Parkinson’s disease and essential tremor, with an estimated occurrence of 0.015-0.06% in the general population (Epidemiological Study of Dystonia in Europe Collaborative, 2000; Steeves et al., 2012). The link between primary dystonias and brain dysfunction, particularly in the basal ganglia, was recognized in the 1970s, thanks to Charles Marsden and colleagues (Berardelli et al., 1998; Marsden et al., 1976). Current understanding suggests dystonic patients experience disrupted cortico-striatal function, leading to impaired striatal control of the GPi, a significant factor in dystonic dysfunction (Berardelli et al., 1998). The proposed mechanism involves disruption and altered synaptic plasticity in the cortico-basal ganglia-thalamo-cortical circuit (Quartarone and Pisani, 2011; Quartarone and Hallett, 2013; Schirinzi et al., 2018; Vitek, 2002), causing irregular neuronal firing patterns (Berardelli et al., 1998) and diminished inhibitory control (Gernert et al., 2000) due to interneuron loss, potentially shifting signalling towards the direct pathway (Wichmann and Dostrovsky, 2011). Within the network, these disturbances manifest as decreased cortical inhibition (Beck and Hallett, 2011; Meunier et al., 2012), persistent β-band synchronization during movement initiation (Crowell et al., 2012), and predominant low-frequency pallidal activity in the α-band at rest (Kuhn et al., 2008). Recent studies indicate lower frequency oscillations, particularly θ and α, are associated with dystonia and can be modulated by DBS (Barow et al., 2014; Knorr et al., 2021; Neumann et al., 2015), while β oscillations appear less significant. Overall, there seems to be widespread reduced inhibitory tone within the extended network (Tisch et al., 2007a), though it remains unclear if it affects the entire network or specific parts (Batla et al., 2015). While cerebellar dysfunction is being discussed as inductor of dystonic symptoms (Brown et al., 2023; Prudente et al., 2014; Schirinzi et al., 2018; Shakkottai et al., 2017), and we indeed recently found electrophysiological evidence for cerebellar dysfunction in the dt^sz^ dystonic hamster model (Kragelund et al., 2025), altered striatal function is likely a primary factor in primary dystonias.

Deep brain stimulation (DBS) has emerged as a promising therapeutic approach for refractory movement disorders, including dystonia (Bledsoe et al., 2020; Krack et al., 2019). By delivering electrical pulses to specific brain regions, DBS modulates abnormal neuronal circuits, restoring more physiologically appropriate activity patterns. The entopeduncular nucleus (EPN), homologous to the human globus pallidus internus (GPi), is a common target for DBS in dystonia, demonstrating significant improvement in motor symptoms. Clinical trials of DBS for dystonia (Volkmann et al., 2012; Volkmann et al., 2014) have shown considerable success, especially in patients with genetic isolated dystonias, mobile dystonias, younger individuals, and those without additional health issues (Bledsoe et al., 2020). However, our understanding of DBS mechanisms in dystonia remains limited, similar to the gaps in understanding dystonia’s pathomechanisms. This limitation arises from several factors: DBS has been primarily studied in Parkinson’s disease (PD) patients and animal models (Udupa and Chen, 2015), offering limited insights into dystonia due to different target nuclei (GPi vs. nucleus subthalamicus). Additionally, most hypotheses are derived from DBS in normal primates, DBS-like stimulation in vitro using normal rodent tissue, or cortical and basal ganglia recordings in patients, restricting our ability to assess the entire network. Unlike most motor symptoms in PD, DBS effects in dystonia require hours of stimulation, suggesting functional network changes (Herrington et al., 2016).

Regarding dystonia, studies show pallidal DBS decreases low-frequency (4–12 Hz) activity, indicating dystonia severity (Scheller et al., 2019), and reduces coherence between pallidal and cortical activity (Barow et al., 2014). Research suggests pallidal DBS in dystonic patients has inhibitory network effects. Increased cortical excitability and synaptic plasticity appear to normalize (Tisch et al., 2007a; Tisch et al., 2007b). Thalamic neuron firing is altered, with most neurons showing reduced firing and a minority showing increased firing (Montgomery, 2006). These findings suggest pallidal DBS diminishes cortical excitability; effects on the broader network remain unknown.

Animal studies have yielded conflicting results. In normal primates, pallidal DBS suppressed neuronal firing through GABAergic afferent activation (Chiken and Nambu, 2013). Conversely, DBS-like stimulation in normal rat brain in vitro caused prolonged afterdepolarizations mediated by cholinergic inputs without neuronal silencing (Kim et al., 2008). Most animal studies have been conducted on healthy controls. In two related dystonia model studies, DBS stimulation was administered under deep urethane anaesthesia (Leblois et al., 2010; Reese et al., 2009), which distorts cortico-striatal connectivity (Paasonen et al., 2018). In a study on a double-hit rat dystonia model (ΔETorA mutation in combination with sciatic nerve injury), DBS attenuated the resting state θ-power reduction otherwise observable in GPi LFP (Knorr et al., 2021) in these animals.

Given this limited knowledge on DBS mechanisms, our group has focused on studying DBS effects in a viable in-vivo dystonia model expressing spontaneous dystonic symptoms: the dt^sz^ hamster model, resembling generalised paroxysmal dystonia in humans, extensively characterised (Perl et al., 2022; Richter and Löscher, 1998). Although generalisability to human primary dystonia is unclear, important similarities exist: This strain shows spontaneous paroxysmal dystonic attacks provoked by handling and stress. As speculated for some human dystonias (Quartarone and Pisani, 2011; Quartarone and Hallett, 2013), this model is associated with increased cortico-striatal excitability (Avchalumov et al., 2013), due to reduced intra-striatal GABAergic signalling, resulting in increased EPN/GPi inhibition (Gernert et al., 2000). Using this model, our group recently reported that short-term DBS dampens synaptic cortico-striatal communication (Heerdegen et al., 2021). Further, we showed that pallidal DBS induced network-wide effects as far as cerebellar activity, normalising firing rates, spike amplitudes, and connectivity measures, with higher connectivity in both healthy and DBS groups compared to dt^sz^ (Kragelund et al., 2025). Altogether, this appears to impact cerebellar nuclei (receiving inhibitory cerebellar projections) as expected, with higher cerebellar activity resulting in dampened activation of these nuclei (Luttig et al., 2024).

An important issue, however, which no study so far has addressed, is whether any adaptive processes take place during DBS. In other words: How does prolonged DBS influence excitability within the motor network? The current study therefore explores these adaptive mechanisms to DBS by comparing short-term and long-term DBS on striatal medium spiny neurons (MSNs) in the dt^sz^ hamster model. By analysing changes in membrane properties, synaptic transmission, and acetylcholine (ACh) modulation, we seek to elucidate the temporal specificity of DBS effects and contribute to optimizing therapeutic strategies for dystonia.

## Methods

### Animal Model

We used the dt^sz^ hamster as a model for paroxysmal dystonia, known for exhibiting atypical neuronal excitability and synaptic activity in the striatum. The study involved 50 adult hamsters, divided into four groups, i.e. animals receiving short-term DBS, long-term DBS, or sham stimulation of either duration. All experimental procedures adhered to institutional guidelines and received ethical approval by the corresponding animal licensing body (Landesamt für Landwirtschaft, Lebensmittelsicherheit und Fischerei Mecklenburg-Vorpommern, licenses number 7221.3-1-053/17 and 7721.3-1-029/20).

### DBS Protocols

DBS was administered to the EPN using implanted bipolar electrodes. Short-term DBS protocol involved a single 3-hour continuous stimulation session, while the long-term protocol consisted continuous stimulation over 11 days (chosen as maximum duration due to progressive remission to be expected after the 45^th^ day of life (Richter and Löscher, 1998), allowing the animal to move freely. The stimulation parameters were set at 130 Hz, 50 μA, and 60 μs, which corresponds to clinical practice and is effective in reducing dystonic symptoms also in our model (Paap et al., 2021). Subjects in the sham group underwent identical surgical procedures but did not receive active stimulation. The specific procedure was as follows:

Animals (32–40 days old) were secured in a stereotactic frame (Narishige, Japan) under deep isoflurane anaesthesia (Isofluran, Baxter, Deerfield, IL, USA; Univentor 1200 Anaesthesia Unit + Univentor 2010 Scavenger Unit, Biomedical Instruments, Zöllnitz, Germany). The periosteum was additionally treated with bupivacaine, a local anaesthetic (bupivacaine 0.25% JENAPHARM®). Two concentric bipolar electrodes (platinum-iridium Pt/Ir; SNEX-100, Microprobes, Gaithersburg, MD, USA) were bilaterally inserted into the entopeduncular nucleus (EPN; equivalent to GPi in humans; stereotaxic coordinates AP: −0.6 mm, ML: ± 2.2 mm, DV: −0.6 mm relative to Bregma from the golden hamster atlas (Wood, 2000). To ensure stable electrode placement, two screws were anchored in the skull behind the electrodes and sealed with dental adhesive (Heliobond + Compo glass flow, Schaan, Liechtenstein; SDR® flow+, Dentsply DeTrey GmbH, Konstanz, Germany). The programmable stimulator (Plocksties et al., 2021)(Institute of Applied Microelectronics, Faculty of Computer Science and Electrical Engineering, University of Rostock) itself was placed in a skin pouch on the back of the hamster. DBS (130 Hz, 50 μA, 60 μs pulse duration) was switched on magnetically after a recovery period of three days; animals remained awake and freely moving. Every second dt^sz^ or control hamster underwent sham stimulation (electrode implantation without active stimulation) to enable comparison of effects with and without stimulation, having confirmed, in a previous study, that sham stimulation has no impact on dystonia severity (Paap et al., 2021). Given that the handling procedure required for staging can alter cortico-striatal network properties (Avchalumov et al., 2013), the animals were sacrificed immediately after DBS.

### Electrophysiological Recordings

Following decapitation, striatal slices were prepared for patch-clamp recordings. The dorsal striatum was sectioned into 300 µm thick brain slices using ice-cold artificial cerebrospinal fluid (aCSF) and maintained at 32°C during recordings, as described previously (Heerdegen et al., 2021). Whole-cell recordings from MSNs were used to assess membrane potential changes, input resistance, and synaptic events, including miniature excitatory postsynaptic currents (mEPSCs). To isolate postsynaptic events and spontaneous synaptic release, mEPSCs were recorded in the presence of tetrodotoxin (1 µM), which blocks action potentials.

### Brain slice preparation for analysis of striatal network excitability and inhibitory tone

Following bilateral DBS or sham stimulation, the animals were euthanized by decapitation while under deep anaesthesia. The electrodes were meticulously extracted from the skull, avoiding any shearing movements, and the brain was swiftly removed and cooled in an ice-cold sucrose solution. This solution comprised (in mM): NaCl 87, NaHCO3 25, KCl 2.5, NaH2PO4 1.25, CaCl2 0.5, MgCl2 7, glucose 10 and sucrose 75. The brain was then dorsally incised at a 40–45° angle to the horizontal plane to preserve cortico-striatal connections (Kawaguchi et al., 1989; Schlösser et al., 1999), and affixed with the cut surface to the microtome platform (VT1200S, Leica Biosystems Nussloch, Germany) (Fig. 1A). The angled brain was horizontally sectioned into para-horizontal slices of 400 μm or 300 μm thickness (for field or patch clamp recordings, respectively), maintaining synaptic connections between motor cortex and striatum. Post-sectioning, the slices (totalling n = 120 from control and n = 174 from dt^sz^ hamsters) were incubated for 60 min in sucrose solution at room temperature. They were then transferred to an interface-type recording chamber (BSC-BU, Harvard Apparatus Inc., March-Hugstetten, USA) perfused with artificial cerebrospinal fluid (ACSF). The ACSF contained (in mM): NaCl 124, NaHCO3 26, KCl 3, NaH2PO4 1.25, CaCl2 2.5 and glucose 10, and was maintained at a constant temperature of 32 °C (TC-10, npi electronic GmbH, Tamm, Germany).

**Figure 1.**
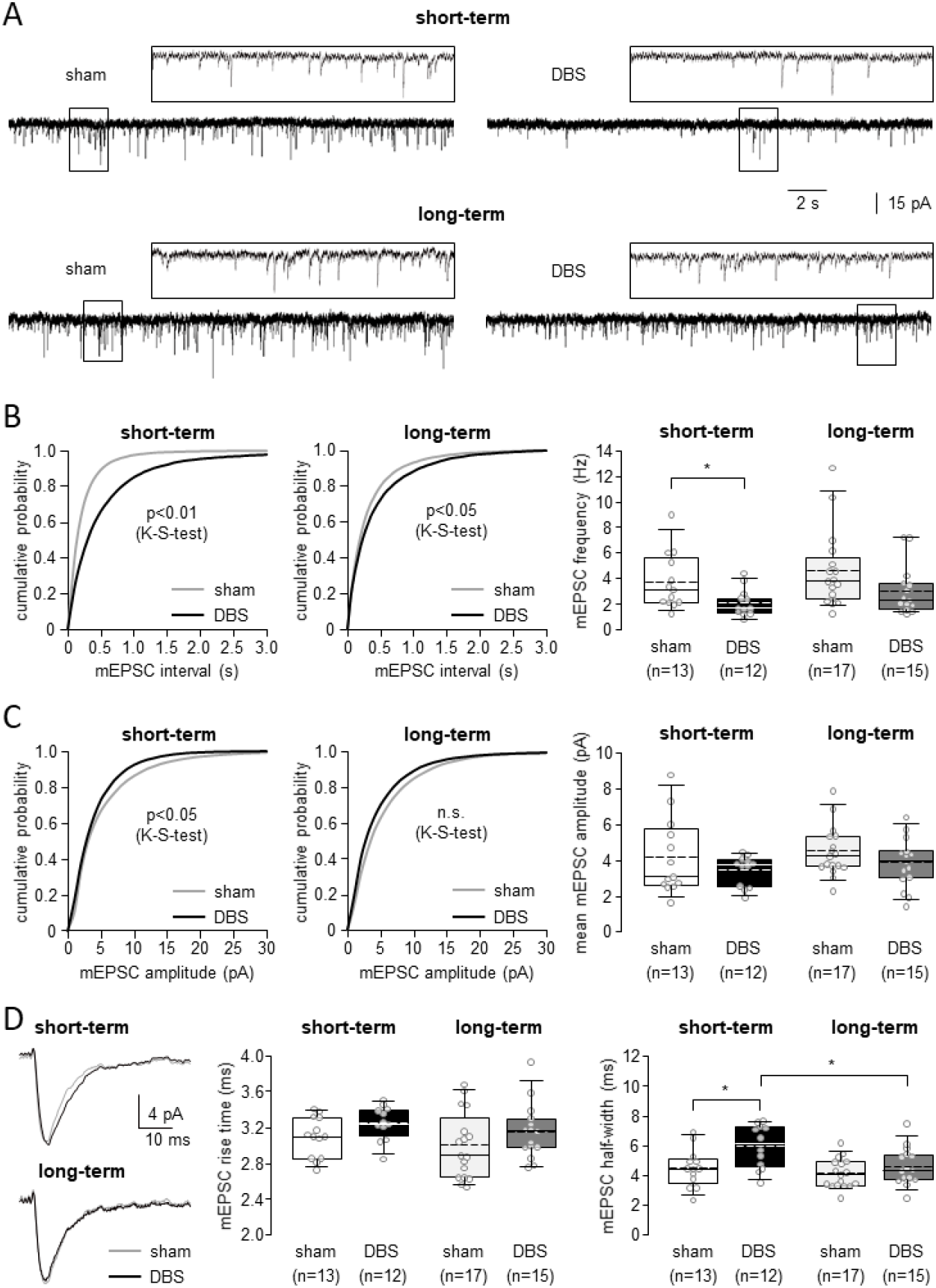
Effect of DBS on miniature EPSCs (mEPSC): *Only short-term DBS reduces the excitability of the synaptic network by decreasing the mean frequency of mEPSCs occurring independently of TTX-sensitive potassium channels at the cortico-striatal synapse of medium spiny neurons*. **(A)** Original traces of mEPSCs recorded from medium spiny neurons in slices from dt^sz^ hamsters subjected to short-term or long-term EPN-DBS (DBS) or sham stimulation (sham), respectively. **(B)** Corresponding cumulative probability histograms and box and whisker plots of the same groups (sham: light curves and boxes, DBS: dark curves and boxes) show that short-term and long-term DBS significantly shifts the interval distribution to the right in dystonic animals. In contrast, only short-term DBS significantly reduces the mean mEPSC frequency (indicated by an asterisk in B, p<0.05, Mann-Whitney rank sum test). **(C)** Cumulative histograms of amplitudes display a significant shift to smaller amplitudes after short-term DBS. The mean amplitudes of mEPSCs are not altered by DBS compared to sham stimulation in either short-term or long-term stimulated dt^sz^ hamsters, as shown by box and whisker plots of mean mEPSC amplitudes. **(D)** Sample traces of mean amplitudes suggest that short-term DBS affects the kinetics of mEPSCs. Box-and-whisker plots demonstrate that neither DBS protocol significantly impacts the rise time of mEPSCs, but the half-width of mEPSCs increases after short-term DBS. In all plots, dark symbols represent data from animals that received EPN DBS, and light symbols represent data from animals that received sham stimulation only. In all box plots, medians are represented by straight lines and means by dashed lines. Single points represent the mean of one experiment (slice).

### Patch-clamp recordings

Medium spiny striatal neurons (MSN) were examined using patch-clamp recordings to evaluate the frequency and kinetic characteristics of spontaneous miniature excitatory postsynaptic currents (mEPSCs). This method was employed to assess presynaptic cortico-striatal functional modifications and to measure MSN properties, which were identified by their distinctive firing patterns. The recordings were conducted at room temperature in cortico-striatal slices immersed in recording ACSF. Borosilicate pipettes (3.1–8.5 MΩ, average 5.6 ± 0.1 MΩ, n = 65, created with DMZ Zeitz puller, Zeitz-Instrumente Vertriebs GmbH, Martinsried, Germany) were filled with a solution comprising (in mM): K-gluconate 115, KCl 20, MgCl2 2, HEPES 10, Na2-ATP 2, Mg-ATP 2, GTP tris 0.3; pH adjusted to 7.3 and osmolarity to 280 ± 5 mosmol/l. MSN were observed using differential interference contrast microscopy and a CCD camera (Till Photonics, Gräfelfing, Germany), allowing visual distinction between MSN and other striatal neurons based on cell shape and size (Siep et al., 2002). Visual classification was further verified through electrophysiological characterization of MSN, revealing specific passive and active membrane properties. The MSN recording seals exceeded 1 GΩ (6.0 ± 1.5 GΩ, n = 65); liquid junction potentials (12.97 mV) and series resistance (18.0 ± 0.7 MΩ, n = 65) were not adjusted for. Voltage- and current-clamp data were acquired using an EPC-10 amplifier (HEKA, Lambrecht, Germany), filtered at 1 kHz, digitized at 20 kHz, and stored using Patchmaster v2.20 software (HEKA, Lambrecht, Germany). The resting membrane potential was initially measured after establishing whole cell configuration. Action potential count, threshold current (rheobase), and the latency of the first action potential at rheobase were determined at 0 pA holding current by applying depolarizing current injections lasting 500 ms, ranging from 0 to at least 300 pA (50 pA increments). Cellular input resistance was derived from the slope of the steady state current– voltage relationship resulting from voltage steps (2 mV increments, 1 s duration) from −60 mV to −80 mV, with a holding potential of −70 mV. For mEPSC measurements, the membrane potential was held at −70 mV, and TTX (1 μM) and gabazine (5 μM) were added to the ACSF. mEPSC events were low-pass filtered at 1 kHz and detected within 5 min with a signal to noise ratio of 5:1 using MiniAnalysis v.6.0.7 software (Synaptosoft, Decatur, USA). The mean mEPSC frequency for each MSN was calculated by dividing the detected events by observation time (300 s). Cumulative probability plots of mEPSC-intervals were generated for each MSN using 50 ms bins. The cumulative plots represent the mean probability of all MSN within the experimental group (for clarity, without indicating SEM). Patch-clamp data were analysed offline using Fitmaster v2.11 software (HEKA), Office Excel 2003 (Microsoft, Redmond, USA) and SigmaPlot 10.0 (Systat Software GmbH, Erkrath, Germany).

### Data Analysis

Prior to commencing the study, sample size calculations were performed using the online Sample Size Calculator (https://clincalc.com/stats/samplesize.aspx). These calculations assumed 20% differences in means, an α-error below 5%, a β error under 20%, and a data variability of 12%. The resulting ideal sample size was determined to be 6 or greater. For experiments involving dystonic animals, which were the primary focus of this study, the sample size was set at 11 or more, while control animal experiments used 6 or more subjects. Signal 2.16 software (Cambridge Electronic Design, Cambridge, UK) was employed to analyse extracellular recording data. All results are presented as means ± SEM, with n representing the number of slices unless otherwise noted. Statistical analyses were conducted using SigmaStat and SigmaPlot software (Systat Software Inc., San Jose, CA, USA). To evaluate the significance of differences in median values of input-output activity between stimulated and sham-stimulated dt^sz^ mutant and control groups, a two-way repeated measures ANOVA (two factor repetition) was used, followed by a post-hoc Holm-Sidak multiple comparison procedure. Cumulative probability plots were statistically analysed using the Kolmogorov–Smirnov test (KS). For all other analyses, the Wilcoxon Rank Sum Test was applied to paired data, while the Wilcoxon-Mann-Whitney Rank Sum Test was used for unpaired data. Statistical significance was set at p < 0.05, denoted by asterisks (* unpaired test) and hash symbols (# paired test). In all figures, whiskers indicate the 5th and 95th percentile limits.

## Results

### Short-term DBS Decreases mEPSC Frequency

We were first interested in a comparison of short- and long-term DBS regarding spontaneous miniature excitatory postsynaptic currents (mEPSC) in medium spiny striatal neurones, which reflect action potential-independent synaptic glutamate release presumably from motor cortical inputs, since we had found these to be reduced in a previous study on short-term DBS, then still using a tethered stimulator (Heerdegen et al., 2021). Also in the current study, short-term DBS, now delivered by a chronically implanted stimulator, resulted in a significant reduction in the frequency of mEPSCs recorded from MSNs compared to sham-treated animals (Fig. 1). Thus, the cumulative probability distributions of inter-event intervals demonstrated a right-ward shift in the DBS-treated group, such that 80% of all intervals were up to 0.87 s long (vs. 0.37 s in sham preparations, Fig 1B, p<0.01), indicating a decrease in the rate of spontaneous excitatory synaptic events (reducing mean mEPSC frequency by about 50%, i.e. from 3.9 Hz for sham vs. 2.0 Hz for short-term DBS, Fig. 1B, p<0.05). This decrease replicates and confirms our initial findings (Heerdegen et al., 2021) of a reduction in the excitatory drive onto MSNs following short-term DBS in a previous set of experiments, potentially through reduced presynaptic glutamate release probability. In contrast, the effect of long-term DBS was somewhat attenuated: While the significant right shift towards longer intervals persisted in reduced form, with 80% of all intervals being up to 0.74 s long vs. 0.52 s in sham preparations (Fig 1B, p<0.05, Fig. 1B), mEPSC frequency relative to sham controls was reduced, but not significantly so (mean frequency: 3.8 Hz for sham vs. 1.6 Hz for long-term DBS; Fig. 1B, n.s.). This seeming lack of effect on frequency can likely be attributed to the fact that mEPSC frequency in two preparations under DBS actually were very high (around 7 Hz, while all other values were below 4 Hz). Amplitudes of mEPSC amplitudes and rise times remained unchanged in any condition (Fig. 1C). There was, however, a difference in mEPSC half-width, which selectively rose in preparations with short-term DBS only (Fig. 1D; 4.6 ms to 6.2 ms, p<0.05) and was thus different from long-term DBS, again reflecting that short-term DBS appears to have more impact on synaptic modulation than long-term. The lack of change in mEPSC frequency after long-term DBS suggests a potential adaptation or homeostatic mechanism that could counteract the initial reduction observed with short-term stimulation. This finding highlights the importance of considering the duration of DBS treatment when evaluating its effects on synaptic transmission.

### Short-term DBS also decreases spontaneous EPSC

In view of these data, we were then interested whether sEPSC, i.e. spontaneous EPSC arising from both, action potential-independent (i.e. mEPSC) and action potential-dependent glutamate release, would be similarly affected. In essence, we indeed see the same results (Fig. 2), perhaps even in a more conclusive way: again, interval lengths were shifted to the right, significantly now exclusively for short-term DBS (80% of all intervals are shorter than 0.41 s in sham, and 0.85 s in DBS preparations, p<0.001, Fig. 2B), which is mirrored in the mean frequency dropping from 4.83 Hz (sham) to 2.06 Hz (DBS), while there is no change in long-term DBS. Again, of all other parameters, only the sEPSC half-width is elevated, too, in short-term DBS (Fig. 2D).

**Figure 2.**
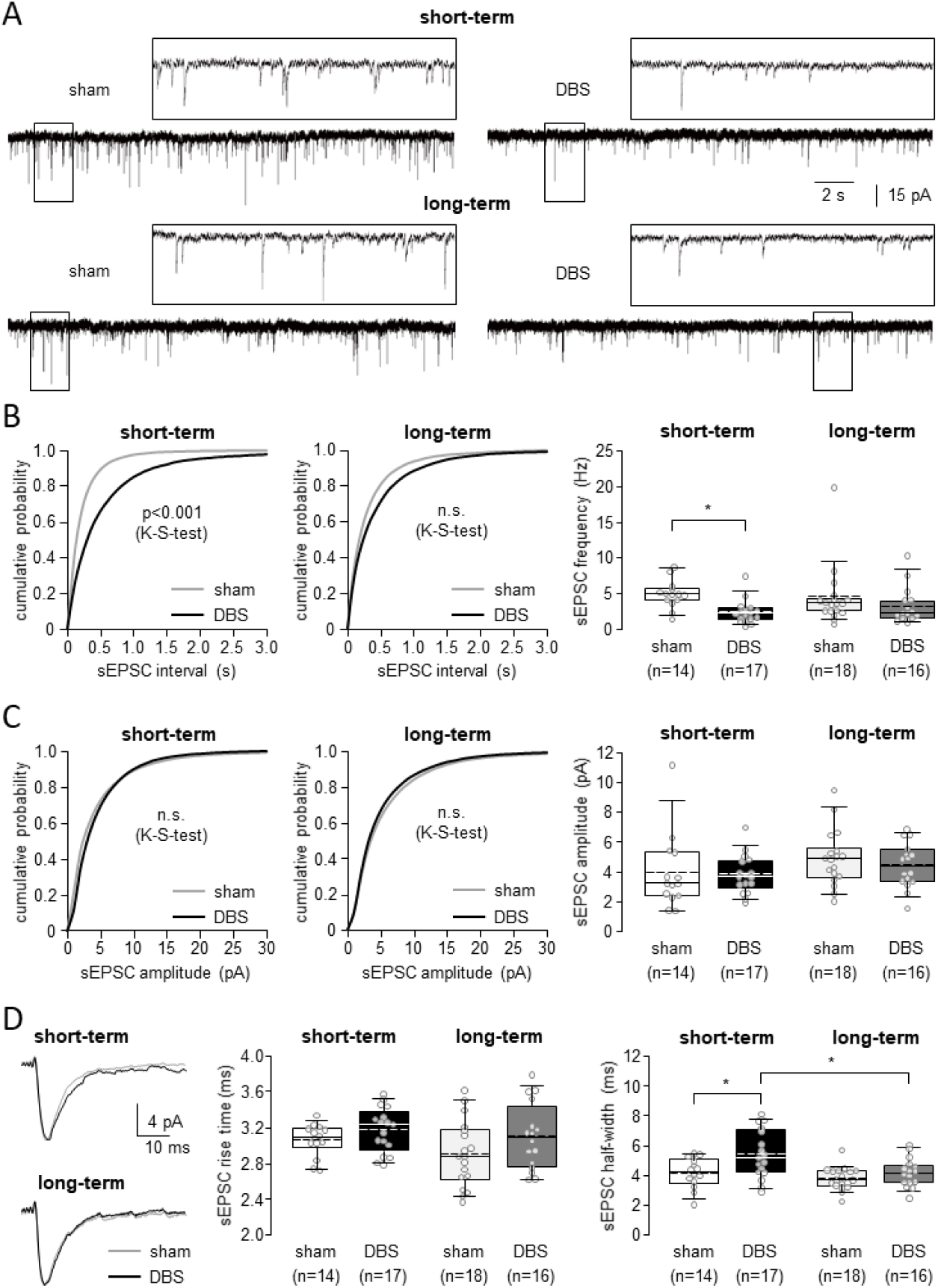
Effect of DBS on spontaneous EPSCs (sEPSC): *Only short-term DBS reduces the excitability of the synaptic network by decreasing the mean frequency of sEPSCs at the cortico-striatal synapse of medium spiny neurons.* (A) Original traces of sEPSCs recorded from medium spiny neurons in slices from dt^sz^ hamsters subjected to short-term or long-term EPN-DBS (DBS) or sham stimulation (sham), respectively. (B) Corresponding cumulative probability histograms and box and whisker plots of the same groups (sham: light curves and boxes, DBS: dark curves and boxes) show that short-term and long-term DBS significantly shifts the interval distribution to the right in dystonic animals. In contrast, only short-term DBS significantly reduces the mean sEPSC frequency (indicated by an asterisk in B, p<0.05, Mann-Whitney rank sum test). (C) The mean amplitudes of sEPSCs are not altered by DBS compared to sham stimulation in either short-term or long-term stimulated dt^sz^ hamsters, as shown by box and whisker plots and cumulative histograms of mean sEPSC amplitudes. (D) Sample traces of mean amplitudes indicate that short-term DBS primarily affects the kinetics of sEPSCs. Box-and-whisker plots reveal that neither DBS protocol significantly impacts the rise time of sEPSCs; however, the half-width of sEPSCs increases with short-term DBS. In all plots, dark symbols represent data from animals that received EPN DBS, while light symbols indicate data from animals that only received sham stimulation. In the box plots, medians are shown as straight lines and mean values as dashed lines. Individual points represent the mean from each experiment. (A) Original traces of membrane potential recordings illustrate how medium spiny neurons in slices from dystonic animals react to EPN-DBS (DBS) or sham stimulation (sham). Firing was triggered by injecting increasing amounts of a depolarising current. The corresponding input-output curve is displayed below the original traces (at 300 pA step current), as a dot diagram (mean ± SEM) of action potential number plotted against current injection. (B) Box and whisker plots of rheobase and of latency to first action potential after depolarising current injection in combination with truncated sample trace show there was, in turn, no effect of short-term and long-term DBS in tissue from dystonic animals on these characteristics. (C) The voltage-current relationship to determine input resistance displays the characteristic rectification of medium spiny neurons in dystonic tissue. Box and whisker plots of input resistance, membrane capacitance and resting membrane potential provide a comparison of data from animals having undergone DBS with data from animals with sham stimulation. In all plots, filled symbols represent data from animals having received short-term DBS, and open symbols represent those of the corresponding animals with sham stimulation only. In all box plots, the median is represented by a straight line and the mean by a dashed one. Single dots represent the arithmetic mean of one experiment (medium spiny neuron).

Comparing mEPSC and sEPSC (action potential-independent only, and action potential-dependent and –independent EPSC), it appears that only in slightly younger dystonic animals (i.e. the 3h-sham DBS group), there is some perceivable contribution of action potential-dependent EPSC, since in this group the frequency of sEPSC was slightly higher than that of mEPSC (3.9 vs. 4.83 Hz, n.s.). Eleven weeks later, mEPSC frequencies rise to ∼ 5Hz, and the action potential-dependent component of EPSC becomes negligible as sEPSC have a similar frequency. DBS, in turn, appears to have a more prominent effect on exactly these action potential-dependent EPSC in the short-term group, since TTX, a sodium channel blocker, only elicits a significant reduction in sham group, but not any more in the DBS group, meaning to say that once DBS reduced EPSC, there is no further reduction possible when sodium channels are blocked (Fig. 3).

**Figure 3.**
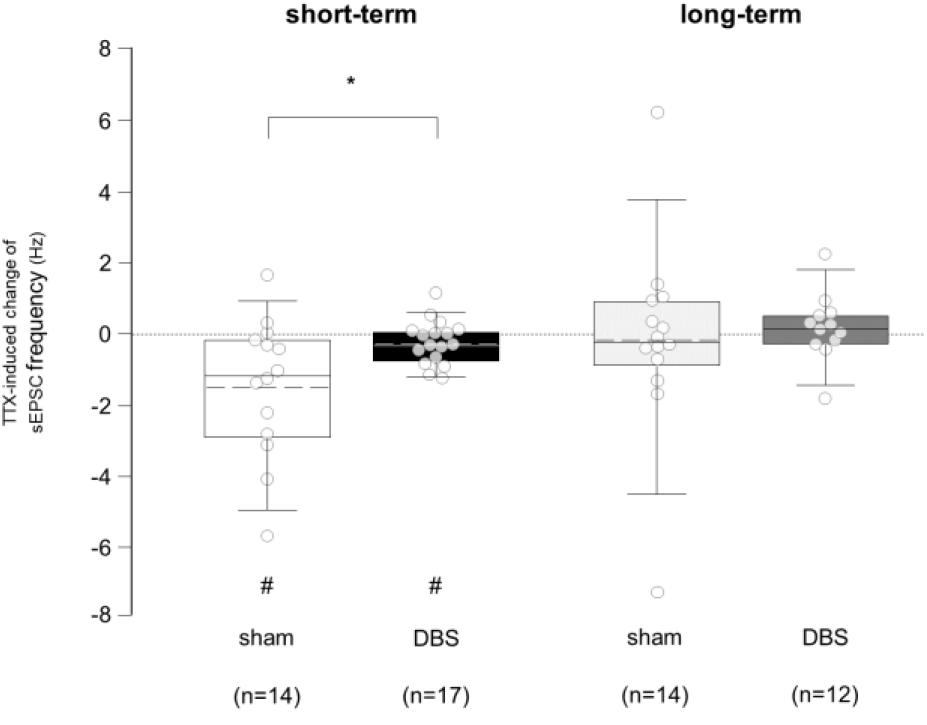
Effect of voltage-gated sodium channels on spontaneous EPSC (sEPSC): *short-term DBS reduces the occurrence particularly of TTX-sensitive sEPSC at the cortico-striatal synapse of medium spiny neurons*. Box and whisker plots show a significant decrease in mean sEPSC amplitude following TTX application in short-term sham-stimulated dt^sz^ hamsters. In contrast, the sEPSC of long-term DBS and sham-stimulated dt^sz^ hamsters remain unaffected by TTX. Dark symbols: data from animals that received EPN DBS. Light symbols: data from animals that only received sham stimulation. In the box plots, medians are shown as straight lines and mean values as dashed lines. Individual points represent the mean from each experiment.

### Effects of DBS on neuronal excitability

As the dampening effect of short-term DBS on synaptic network excitability wanes with prolonged DBS, we were all the more interested in the response of cellular excitability to DBS, and its temporal development. To test this, we elicited action potential firing with increasing current injection, to obtain input-output curves of neuronal firing (Fig. 3). Importantly, deep brain stimulation (DBS) of the dt^sz^ hamster, both short-term and long-term, influences the excitability of striatal medium spiny neurons in contrasting ways: Short-term DBS enhances the cellular action potential firing response to depolarisation, while long-term DBS diminishes it (p<0.01, ANOVA). Thus, long-term DBS apparently reduces excitability on the cellular side, and less so on the synaptic.

### Effects of DBS on passive membrane properties of MSNs

Looking at passive cellular properties, voltage-current (V-I) relationship analysis revealed characteristic rectification properties of MSNs in dystonic animals (Fig. 3C). While there were slight differences between short- and long-term DBS cells (more rectification in short-term, as well as slightly increased resting membrane potential) overall, none of the passive properties rheobase, latency to first AP, input resistance, capacitance and resting membrane potential) differed between the two groups (Fig. 3B,C), confirming the changes in firing rates are likely to be related to the properties of e.g. sodium channels rather than to passive membrane characteristics.

### Acetylcholine (ACh) Modulation of Synaptic Transmission

Cholinergic neurons in the striatum are very prominent and widely interconnected interneurones (Pisani et al., 2007). Specifically, they are thought to raise the efficacy of cortico-striatal signalling (Deffains and Bergman, 2015), and are thus excellently poised to modulate synaptic tone (Fino et al., 2008; Fino et al., 2010). We were therefore interested in how this type of modulatory effect would also be influenced by DBS, and tested this by exogenous application of ACh. Indeed, this led to a differential response in mEPSC frequency between short-term and long-term DBS groups (Fig. 3). In the short-term DBS group, the application of ACh resulted in a significant increase in the frequency of miniature excitatory postsynaptic currents (mEPSCs) between nerve cells.

Specifically, the frequency rose from 1.6 to 2.4 Hz (p < 0.05), representing a substantial enhancement in synaptic activity. This observation suggests that in the early stages of DBS treatment, cortico-striatal communication shows increased responsiveness to ACh, on the basis of an overall reduced spontaneous synaptic output (see Fig. 1). This interaction is complex, though, as ACh actually reduces the amplitude of mEPSC after short-term DBS (Fig. 4B), and incidentally decay times of these mEPSC in both groups, short- and long-term DBS (Fig. 4C). Conversely, the long-term DBS group exhibited a markedly different response to ACh application: In this cohort, the same administration of ACh produced only an attenuated effect on mEPSC frequency, with a slight increase from 2.2 to 2.3 Hz (p > 0.1). This negligible change indicates a significant reduction in cholinergic modulatory effect on cortico-striatal connections over extended periods of DBS treatment, and suggests an attenuation adaptation of cholinergic modulation over time, possibly via receptor or signalling downregulation. Importantly, this adaptation again, like the attenuation in synaptic modulatory response in general, favours cellular excitability impact of DBS over synaptic network modulation with prolonged DBS, suggesting that the brain undergoes adaptive changes in response to prolonged stimulation.

**Figure 4.**
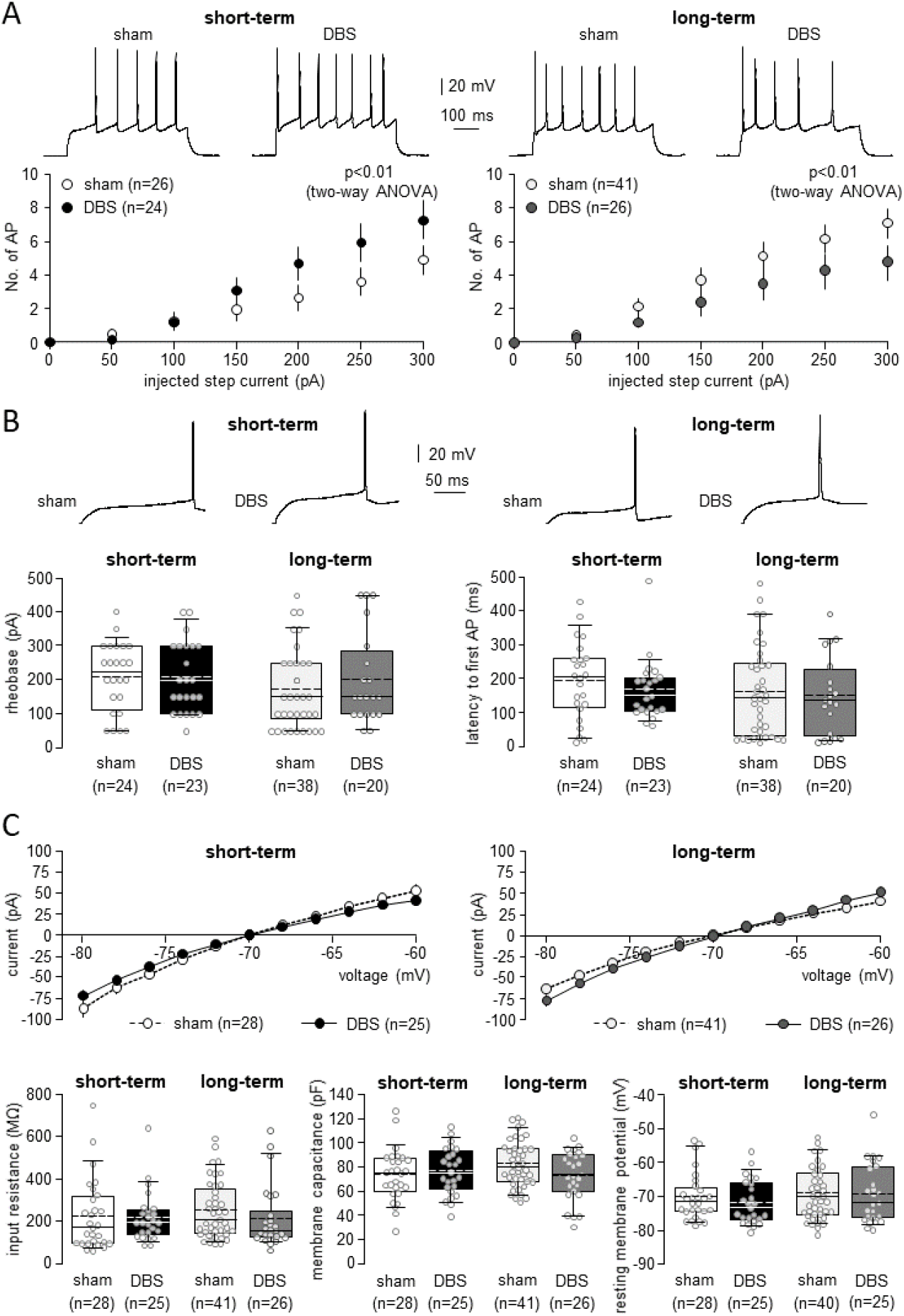
Effect of DBS on cellular excitability: *Short-term and long-term DBS of the dt*^*sz*^ *hamster affects the excitability of medium spiny neurons in the striatum in opposite ways, with short-term DBS increasing cellular action potential firing response to depolarisation, and long-term DBS decreasing it*. (A) Original traces of membrane potential recordings illustrate how medium spiny neurons in slices from dystonic animals react to EPN-DBS (DBS) or sham stimulation (sham). Firing was triggered by injecting increasing amounts of a depolarising current. The corresponding input-output curve is displayed below the original traces (at 300 pA step current), as a dot diagram (mean ± SEM) of action potential number plotted against current injection. (B) Box and whisker plots of rheobase and of latency to first action potential after depolarising current injection in combination with truncated sample trace show there was, in turn, no effect of short-term and long-term DBS in tissue from dystonic animals on these characteristics. (C) The voltage-current relationship to determine input resistance displays the characteristic rectification of medium spiny neurons in dystonic tissue. Box and whisker plots of input resistance, membrane capacitance and resting membrane potential provide a comparison of data from animals having undergone DBS with data from animals with sham stimulation. In all plots, filled symbols represent data from animals having received short-term DBS, and open symbols represent those of the corresponding animals with sham stimulation only. In all box plots, the median is represented by a straight line and the mean by a dashed one. Single dots represent the arithmetic mean of one experiment (medium spiny neuron).

**Figure 5.**
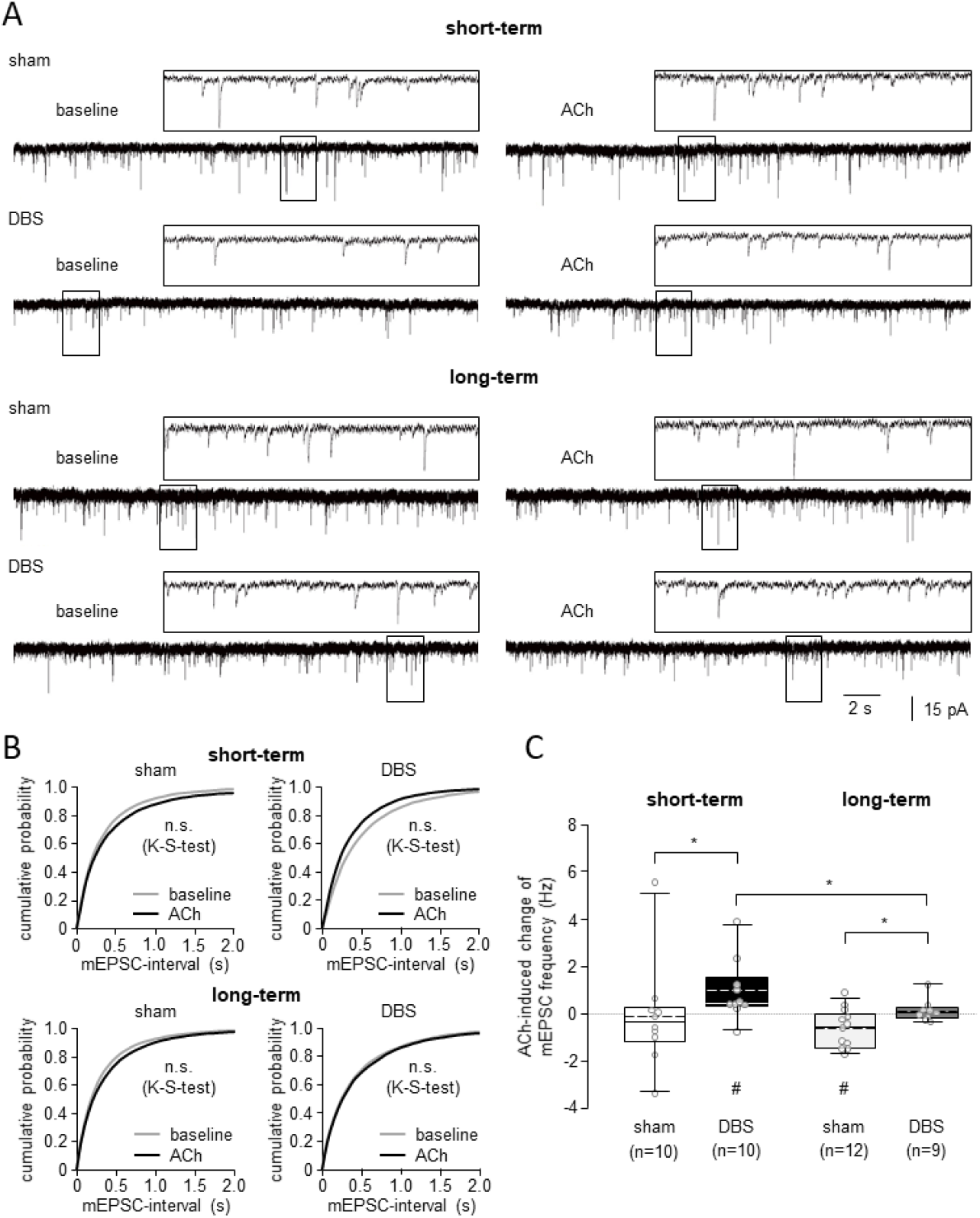
Effect of acetylcholine (ACh) on the frequency of miniature EPSCs (mEPSCs): *Short-term DBS is more potent than long-term DBS in enhancing ACh-induced positive modulation of mean frequency of mEPSCs, which occur spontaneously at the cortico-striatal synapse of medium spiny neurons*. **(A)** Original traces of mEPSCs recorded from medium spiny neurons in slices from dtsz hamsters subjected to short-term or long-term EPN-DBS (DBS) and sham stimulation (sham), respectively. **(B)** Corresponding cumulative probability histograms of the same groups (sham: light curves, DBS: dark curves) show no significant effect of ACh on the interval distribution of mEPSCs.**(C)** In contrast, ACh-induced changes in mean mEPSC frequency differ between neurons from sham-stimulated and DBS-stimulated animals in both short-term and long-term stimulation, but more strongly so in short-term DBS. In all plots, filled symbols represent data from animals that received EPN-DBS and open symbols represent data from animals that received sham stimulation only. In all box plots, medians are represented by straight lines and means by dashed lines. Single points represent the mean of one experiment (slice).

**Figure 6.**
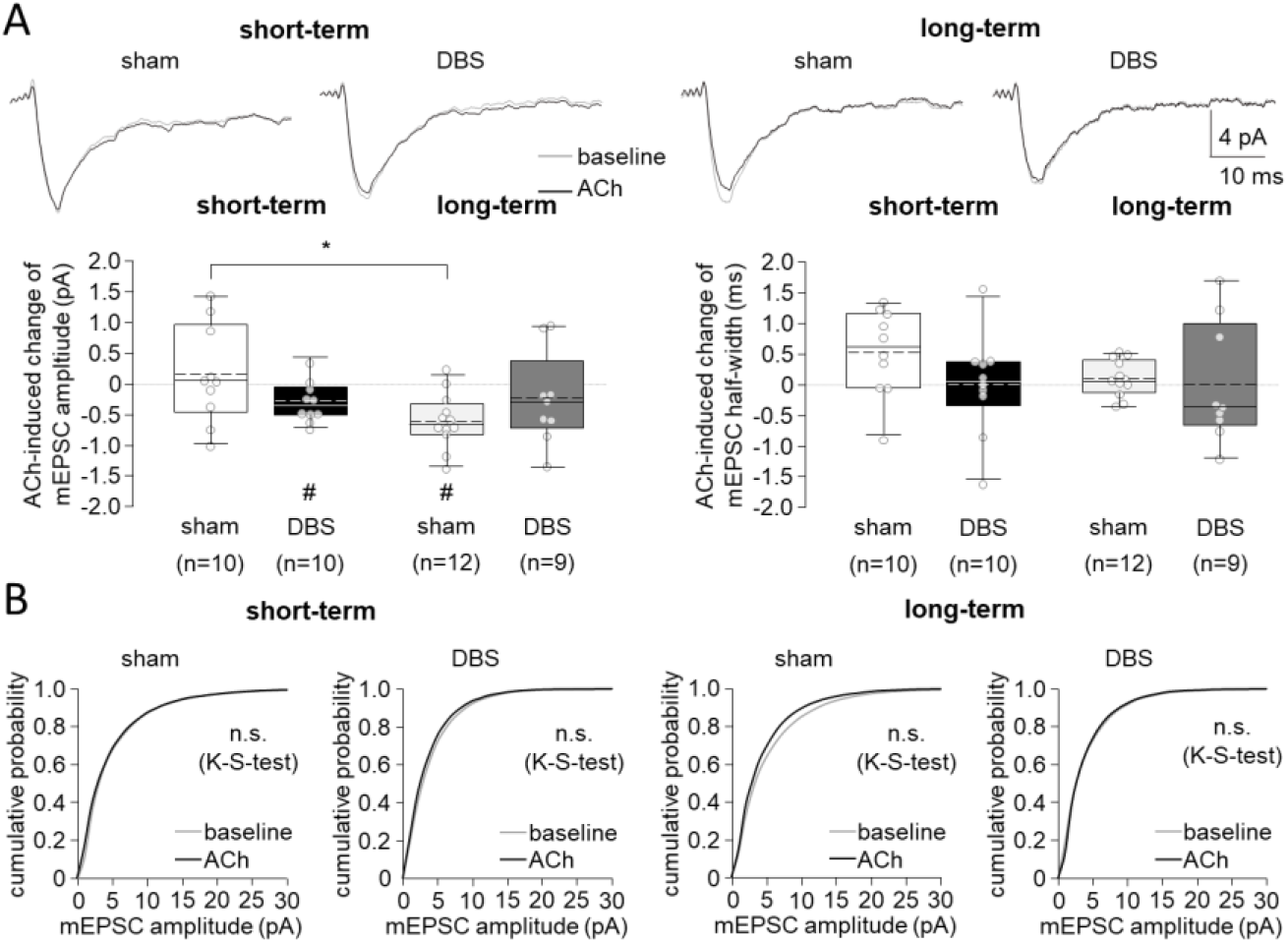
Effect of acetylcholine (ACh) on the amplitude of miniature EPSCs (mEPSCs) *Exogenous ACh has no effect on the amplitude of mEPSC in either tissue of sham stimulated animals or animals with DBS, but differentially modulates the shape of mEPSC, with smaller amlitudes of mEPSC in short-term DBS, and similarly reduced decay times in both, short- and long-term DBS*. **(A)** Sample traces of mean mEPSC amplitudes from individual medium spiny neurons, along with box and whisker plots, reveal a significant reduction in amplitude among hamsters that underwent short-term deep brain stimulation (DBS) and long-term sham stimulation. In contrast, the half-width of mEPSC amplitudes shows no significant changes due to ACh. **(B)** Cumulative probability histograms demonstrate that ACh has no significant effect on the distribution of mEPSC amplitudes in either experimental group.

### Functional impact

To assess overall striatal network output, as a very first approximation, we gauged output by considering the relationship between synaptic input (E) and neuronal excitability reflected in firing rates (F). Output R was then simplistically defined as

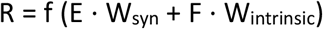

Where E is the excitatory input, W_syn_ and W_intrinsic_ are scaling factors (with W_syn_ + W_intrinsic_ = 2) that represent the relative contribution of synaptic input and intrinsic firing to the network output, and F is the firing rate of MSNs, which changes over time with short-term and long-term DBS effects.

Taking up our findings, we further include the changes in synaptic and intrinsic excitability as follows:

Short-term DBS:

- E (synaptic input) = 1 (baseline) * 0.5 (50% decrease),
- F (firing rate) = 1 (baseline) * 1.5 (50% increase),

Long-term DBS:

- E = 1 (no change)
- F = 1 (baseline) * 0.7 (30% decrease)

Depending on the relative synaptic W_syn_ and intrinsic excitability W_intrinsic_ scaling factors, which we assumed to be either balanced (W_syn_/W_intrinsic_ = 1/1), with increased weight on intrinsic excitability (W_syn_/W_intrinsic_ = 0.8/1.2), or a more important role of synaptic activity (W_syn_/W_intrinsic_ = 1.2/0.8), we thus yield the following values for striatal output, with 2.0 (no cholinergic modulation) and 2.24 (with cholinergic modulation) being the outputs to be expected without DBS:

**Tab. 1.** Striatal output value as sum of excitability and synaptic input (both set to unity at no DBS conditions) at varying degrees of relative weight. The different colours denote differences vs. sham conditions: black: no change, green: reduction, red: increase.

| | cholinergic modulation | $W_{\text{syn}} / W_{\text{intrinsic}}$<br>1 / 1 | $W_{\text{syn}} / W_{\text{intrinsic}}$<br>0.8 / 1.2 | $W_{\text{syn}} / W_{\text{intrinsic}}$<br>1.2 / 0.8 |
| --- | --- | --- | --- | --- |
| Short-term DBS | — | **2.00** | **2.20** | **1.80** |
|  | + | **2.10** | **2.28** | **1.92** |
| Long-term DBS | — | **1.70** | **1.64** | **1.76** |
|  | + | **1.80** | **1.72** | **1.88** |

As can be derived from this table, using our very simple approximation, a reduction of striatal output is apparently best achieved when synaptic input has a 50% higher impact on overall output (W_syn_ = 1.2) than intrinsic excitability (W_intrinsic_ = 0.8). Under this condition, striatal output would be reduced by 14-19%, and could still be modulated by cholinergic input by about 7%. Obviously, these are assumptions which await testing in further experimental and very much refined modelling approaches.

## Discussion

### Short-vs. long-term DBS: Effects on cellular and synaptic excitability

Our investigation reveals that short-term and long-term deep brain stimulation (DBS) elicit contrasting effects on synaptic transmission and neuronal excitability in the dt^sz^ hamster model of dystonia. Short-term DBS induced a substantial decrease in miniature excitatory postsynaptic current (mEPSC) frequency, replicating and confirming our previous findings on short-term DBS using an external stimulator (Heerdegen et al., 2021), and in turn modified the intrinsic membrane properties of medium spiny neurons (MSNs), overall leading to a decrease in network, and an increase in cellular excitability. By contrast, long-term DBS, while continuing to prolong mEPSC intervals, was overall less effective in down-regulating network excitability, with at least some preparations (2/15, see Fig. 1 B) showing little effect on DBS, perhaps reflecting a kind of non-responder phenotype, although such non-reponders were not found in parallel in-vivo experiments assessing dystonia severity after long-term DBS, which proved to be effective (personal communication Angelika Richter). Thus, we have to conclude that the synaptic depressive effect is perhaps less prominent after long-term DBS. Instead, there was a reduction in intrinsic excitability of MSNs, possibly as a homeostatic adaptation to prevent excessive neuronal firing. Importantly, thus, there is a continuing adaptation to long-term DBS, shifting the modulatory impact from synaptic at early stages, to intrinsic excitability changes at late ones.

### Contribution of cellular vs. synaptic changes to dystonic phenotype

As our main finding touches upon the different roles of cellular vs. synaptic changes in dystonia, it is important to re-consider available data (all, obviously, in animal models) on their respective contributions to the dystonic phenotype. Interestingly, regardless of the model, in most instances, synaptic rather than intrinsic changes were documented. Thus, in our own studies on dt^sz^ hamsters, we saw increased synaptic, glutamatergic responses, associated with higher release-probability, and also increased long-term potentiation (Kohling et al., 2004). This is corroborated by findings of increased mEPSC frequency in ΔE-torsinA mutation-affected neurones (a mutation associated with DYT1 dystonia) (Kakazu et al., 2012), and likewise altered synaptic plasticity in a mouse DYT-knock-in model (Martella et al., 2009; Martella et al., 2014). Only under pharmacological challenge (using a sodium-channel blocking drug), evidence can be found for intrinsic, sodium-current changes in dystonic striatal neurones, which were more resistant to blockade (Siep et al., 2002).

### Mechanistic underpinnings of DBS Effects

Considering the effects we describe in the current study, i.e. changes in cortico-striatal synaptic communication (hence changes, very probably, in cortical neurones and their axonal processes / release machinery), and in intrinsic properties of striatal neurones, the question is via which pathway(s) this occurs. The structure being stimulated is the GPi, and thus DBS must be able to influence the motor network to a much farther extent than just the direct stimulation target, a hypothesis which is now being widely accepted for DBS in general (Franz et al., 2023; Gubellini et al., 2009; McIntyre and Hahn, 2010). Via which pathway then will a focal stimulation of GPi impact motor cortex or striatum? Three possibilities readily come into mind – either via the GPi-thalamus-cortex loop, retrogradely via antidromic firing of respective afferents (GPi – striatum, and then striatum – cortex) or by direct effects on the hyperdirect pathway. The latter runs close to the GPi and is likely activated at supra-threshold levels (Franz et al., 2023; Spiliotis et al., 2024) and would at least impact cortical neurones, and then indirectly, again, striatal ones. The signalling pathways or mechanisms responsible for the observed changes mEPSC generation on the one hand, and action potential frequency on the other are unclear. It seems plausible that release probability is affected as underlying cause for mEPSC interval reduction and frequency increase, perhaps in a similar fashion to the findings in ΔE-torsinA-mutated neurons, where vesicular release was increased, without changes in quantal content (Kakazu et al., 2012), even though these were hippocampal neurones, and not striatal ones, and hence the comparability is somewhat limited (and not assessable regarding functional consequences). Nevertheless, in view of the apparent up-regulation of cortico-striatal glutamatergic communication in dystonic conditions (Heerdegen et al., 2021; Martella et al., 2014), this would directly counteract one of the main affected functions. Regarding the apparent shift towards reducing cellular excitability as DBS becomes chronic, one hint comes from in-vitro investigations of subthalamic neurones, showing that these cells can be driven into pronounced slow inactivation of resurgent persistent sodium current components with high-frequency stimulation simulating DBS (Do and Bean, 2003), a current which is also present in striatal neurons (Patel et al., 2016). This might be a possible mechanism, although it would not explain why this effect appears only with long-term DBS, since the cited reports saw changes within minutes. Obviously, elucidating the exact mechanisms will have to be left to future studies.

### Net Change in Basal Ganglia Network Activity

Short-term DBS effects: The reduction in EPSC frequency thus likely results from synaptic plasticity or network feedback in response to DBS. However, with a 50% increase in MSN firing rate, the network output impact is not straightforwardly additive. Despite reduced input, MSNs are more responsive to remaining excitatory input from cortical and pallidal sources, perhaps even increasing striatum output, with direct inhibition of the GPe, and resulting thalamic facilitation in the direct pathway.

Long-term DBS effects: The overall unchanged EPSC frequency and duration suggests synaptic homeostasis over time, involving synaptic scaling or metaplasticity, possibly stabilizing the striatal network against short-term DBS over-excitation, as a compensation for the initial disturbance. Nevertheless, there the now reduced firing rate medium spiny neurones, likely due to intrinsic synaptic or cellular adaptation involving ion channel regulation, should lead to an overall striatum output decreases, reducing network excitability and potentially preventing overactivity leading to motor dysfunctions like dystonia or hyperkinetic movements..

### Functional impact

The most important question is: What is the functional impact of the described changes, and specifically of the continuing adaptation DBS? From the point of network efficiency, we hypothesize that DBS-induced changes to the cortico-striatal network, especially in long-term adaptation, may be influenced by the principle of criticality. Theoretical models suggest that neural networks function most efficiently when they are near criticality—this is when the system has the widest dynamic range for processing inputs (Gautam et al., 2015; Viana et al., 2014). This state, characterized by scale-free fluctuations and optimal balance between excitatory and inhibitory dynamics, supports both local and long-range coupling of oscillatory activity across brain regions, as demonstrated by studies on brain networks at criticality (Avramiea et al., 2022). In the context of DBS, short-term effects involve enhanced intrinsic excitability in medium spiny neurons, which might compensate for decreased synaptic input (Gautam et al., 2015), as would actually also the increase in ACh-mediated upregulation of mEPSP (see Fig. 3). This enhanced excitability could be seen as a temporary shift towards criticality, where the neuronal network becomes more responsive to inputs and exhibits increased sensitivity to synaptic activity (Protachevicz et al., 2018). Over the long term, as the system adapts to DBS, intrinsic excitability of MSNs may downregulate, stabilizing the network to prevent hyperactivity and maintain functionality. This process mirrors how networks at criticality modulate their activity to maintain a functional dynamic range (Avramiea et al., 2022; Protachevicz et al., 2018). This dynamic, where short-term DBS elevates neuronal excitability and long-term DBS stabilizes it through homeostatic mechanisms, could be beneficial for normalizing motor function in basal ganglia disorders. The network adapts by reducing the risk of excessive neural activity—an issue that may cause dyskinesias—and ensures the system does not fall into a hypoactive state (Yousuf et al., 2020). Thus, the modulation of intrinsic excitability through DBS may align with the brain’s natural tendency to maintain criticality, and to choose modulation of intrinsic excitability over critical dampening of synaptic input, since the latter is necessary for optimizing long-range coupling and synaptic communication across the cortico-striatal pathway (Avramiea et al., 2022; Yousuf et al., 2020), which also our modelling studies suggest (Spiliotis et al., 2024) is necessary for basal-ganglia-thalamic coupling in normal motor function.

## Notes

### Competing Interest Statement

The authors have declared no competing interest.

### Summary of Updates

Figure legend of figure 2, which was unfortunately the same as of figure 4 in the previous version

